# Z-Hunt-DP: accelerating thermodynamic Z-DNA prediction with dynamic programming

**DOI:** 10.64898/2026.09.14.751448

**Authors:** Maysara Al Jumaily, Hamza Qureshi, Hongbin Yan, Yifeng Li

**Affiliations:** Department of Computer Science, Brock University, St. Catharines, Ontario, Canada; Department of Chemistry and Centre for Biotechnology, Brock University, St. Catharines, Ontario, Canada; Department of Biological Sciences, Brock University, St. Catharines, Ontario, Canada

## Abstract

Z-DNA is a left-handed DNA conformation implicated in gene regulation and chromatin dynamics. Because it is usually less thermodynamically favorable than canonical B-DNA under physiological conditions, computational tools are needed to identify sequences likely to adopt the Z conformation. Legacy Z-Hunt uses a dinucleotide thermodynamic model, but searches every anti/syn assignment in a window, causing its conformation search to grow exponentially with window size. We present Z-Hunt-DP, an exact dynamic programming reformulation that preserves the original thermodynamic objective while reducing this search from *O*(2^*d*^) to *O*(*d*) for a window of *d* dinucleotide positions. On benchmark windows, Z-Hunt-DP matched the brute-force minimum energy within numerical tolerance and achieved a 3.30 × 10^5^ speedup at 20 dinucleotides. In an interval-localization benchmark on public human loci with experimentally mapped Z-DNA, it recovered the clipped reference interval in all 24 cases and was the most stable localizer under midpoint-core and expanded-panel analyses. Since the comparison set mixes thermodynamic, heuristic, and learned models, these cross-tool results are interpreted as localization comparisons rather than direct thermodynamic score tests. The source code and benchmark materials are available at https://github.com/Aljumaily/Z-Hunt-DP.

**Author summary:** DNA can form structures other than its familiar right-handed double helix. One alternative, Z-DNA, is left-handed and may influence how genes are regulated, but identifying it across long DNA sequences remains computationally demanding. Existing Z-Hunt software estimates Z-DNA propensity from experimentally derived energy values. Its central calculation tests every possible local DNA conformation, so its run time rises very quickly as the sequence window grows. We developed Z-Hunt-DP, which solves the same thermodynamic problem by reusing the best partial solutions at each position. This yields the same optimal energy as exhaustive search but requires far less computation. We confirmed exact agreement with brute-force calculations and found large speed gains on increasing window sizes. We also tested whether predicted regions align with public human loci where Z-DNA has been experimentally mapped. Z-Hunt-DP consistently located the center of those loci more reliably than the other tested tools. The open-source implementation makes this established thermodynamic model practical for reproducible genome-scale analysis and for prioritizing compact experimental validation targets.

## Introduction

DNA is a structurally polymorphic molecule, capable of adopting multiple helical conformations based on its sequence, or environmental conditions such as hydration and ionic composition. Although many conformations exist, the three most well-characterized and biologically relevant forms are A-, B-, and Z-DNA. These differ in helical geometry, groove width, and backbone conformation [1]. The predominant form, B-DNA, exists under physiological conditions and is a right-handed helix with approx. 10 base pairs per turn. A-DNA is also right handed and forms under reduced hydration conditions [2].

In 1972, salt-dependent inversion of the circular dichroism spectrum of poly(dG-dC) was observed [3, 4] and later linked to the left-handed helix now called Z-DNA [4]. Once viewed largely as a biochemical curiosity [5], Z-DNA is now supported by in vivo evidence and implicated in gene regulation and related chromatin processes [6–9]. Additionally, the left-handed conformation of Z-DNA has been shown to exhibit antigenic properties generated by its distinct antigenic surface relative to B-DNA, enabling selective recognition by Z-DNA-binding proteins such as antibodies. Anti-Z-DNA antibodies have been found in systemic lupus erythematosus, an autoimmune disease [10]. More broadly, it illustrates that nucleic acid conformation is part of genetic regulation [11, 12].

At the structural level, Z-DNA differs from B-DNA in its unique zig-zag sugar-phosphate backbone shape and *syn* purines [4, 13]. This shape results from the orientation nucleobases adopt along the bond connecting the deoxyribose to the nucleobase, known as a glycosidic bond. All four nucleobases exist in the *anti* conformation in B-DNA; in contrast, purine bases (guanine (G) and adenine (A)) adopt a *syn* conformation, where the base is rotated such that its six-membered ring lies over the deoxyribose sugar. Furthermore, alternating purine-pyrimidine tracts, especially GC or CA repeats, lower the energetic cost of the B-to-Z transition [13, 14], whose kinetics also depend on ionic conditions and segment length [15]. The *syn* conformation of purines is intrinsically destabilizing due to steric strain arising from base-sugar proximity, and the zig-zag backbone geometry increases the proximity between negatively charged phosphate groups. B-DNA is more stable under low ionic strength; however, high-ionic-strength environments reverse their relative stabilities due to cation-mediated electrostatic screening. Since replication, transcription, and chromatin remodeling alter DNA twist, predicting Z-DNA-prone regions remains important.

The original Z-Hunt addressed this problem with an energy-based framework that evaluates dinucleotide steps and ranks windows against a random-sequence background [16]. Its main computational cost is conformation search: with *d* dinucleotide positions and two local states corresponding to the orientation of each nucleobase along its glycosidic bond within a dinucleotide unit, anti-syn (`AS`) and syn-anti (`SA`). A straightforward then solver checks 2^*d*^ assignments to find the minimum-energy path.

We present Z-Hunt-DP, an open Python implementation built around an exact dynamic programming (DP) reformulation of that search. It utilizes the CPU along with multithreading and without the usage of GPU. Since the energy at each position depends only on the current dinucleotide and the immediately preceding state, the minimum-energy assignment can be solved in *O*(*d*) time without changing the thermodynamic objective. Our contributions are an exact same-objective solver for Z-Hunt, a readable open implementation, and a reproducible benchmark suite that separates brute-force validation from broader interval-localization comparisons. Brute-force and legacy Z-Hunt support the exactness claim; external tools support only the practical localization claim.

## Background

### Thermodynamic basis of Z-Hunt

Z-Hunt employs a sliding window of a user-defined range to divide a genomic sequence into overlapping segments of up to 24 base pairs (bp), and scores each sliding window by the ease of the B-to-Z transition [16]. Each dinucleotide position is assigned one of two states, `AS`or `SA`, yielding four possible transitions between neighbors. For a given window, the algorithm finds the lowest-energy state assignment, computes Boltzmann-weighted quantities, and derives a linking-difference statistic. Ho et al. calibrated these scores against random-sequence backgrounds, so that windows of different lengths and composition could be compared on a common scale [16]. The bottleneck is the state search. Brute-force evaluates 2^*d*^ assignments for *d* dinucleotide positions. Since the cost at position *t* depends only on the states at *t* − 1 and *t* and the dinucleotide at *t*, DP applies directly.

### Current tools and related resources

Later resources use different model families. Experimentally, Li et al. [17] used the Z*α* domain of ADAR1 (Adenosine Deaminase Acting on RNA 1) to map stable Z-DNA segments in the human genome, underscoring that computational predictions must ultimately be checked against biological data. On the software side, Z-DNA Hunter is motif-based [18], DeepZ and Z-DNABERT are learned predictors [19, 20], and ZSeeker uses experimentally tuned sequence weights plus a maximum-subarray scan [21]. Only Z-Hunt and Z-Hunt-DP share the same dinucleotide-energy objective and downstream Boltzmann and linking-difference calculations, so legacy Z-Hunt is the exact same-objective benchmark. Since the public DeepZ release provides published hg19 outputs but no packaged inference route for arbitrary loci, we treat it as an annotation-backed reproducibility check after hg18-to-hg19 liftover. Table 1 summarizes the defensible comparisons across this heterogeneous set.

**Table 1.** Defensible quantitative comparisons across the tools studied.

| Tool | Model family | Typical outputs | Quantitative comparisons |
| --- | --- | --- | --- |
| Z-Hunt [16] | Thermodynamic model with exhaustive anti/syn search | Linking difference, slope, final Z-score, and optimal anti/syn assignment | Direct score agreement, MAE or RMSE on shared outputs, Pearson or Spearman correlation, exact state-string agreement, and runtime on identical windows |
| Z-Hunt-DP (this work) | Thermodynamic model with dynamic programming anti/syn search | Same outputs as Z-Hunt, plus summary tables, region annotations, and publication-ready plots | Exact score and state-string agreement with Z-Hunt, runtime comparison on identical windows, and interval overlap with experimental regions |
| Z-DNA Hunter [18] | Pattern-based motif detector | Interval coordinates, length, GC richness, GT richness, raw score, and score percentage | Interval overlap, covered-base Jaccard index, density per kb, top-hit concordance, and rank correlation only after score normalisation |
| DeepZ [19] | Supervised sequence + feature deep-learning model | Per-base probabilities, hg19 interval annotations, and locus plots | Interval overlap after genome harmonization and threshold calibration; annotation-backed comparison rather than direct thermodynamic score subtraction |
| Z-DNABERT [20] | Transformer fine-tuned on experimental Z-flipons | Region probabilities, genome tracks, attention maps, and mutagenesis heatmaps | Precision or recall, AUROC/AUPRC, interval overlap with experimental regions, and calibrated rank correlation on matched loci rather than raw score subtraction |
| ZSeeker [21] | Experimentally tuned scoring model with maximum-subarray search | Thresholded candidate subsequences, Z-scores, penalty-aware rankings, and downloadable plots | Sequence-level ranking agreement, interval overlap, density per kb, and correlation after score normalisation rather than direct thermodynamic error |

## Materials and methods

### Problem formulation

The algorithm scans DNA in dinucleotide positions because the Z-Hunt energy tables are indexed by consecutive base pairs. Let *x*_*t*_ be the encoded dinucleotide at position *t* and let *s*_*t*_ ∈ {AS, SA} be the state at that position. For the canonical bases in Table 2, we will encode each base as a two-bit digit, where A → 00, C → 01, G → 10, and T → 11. In case *b*_*t*_ and *b*_*t*+1_ are adjacent base codes, then

**Table 2.** Canonical Z-Hunt coefficients for the standard A/C/G/T bases.

| Transition | AA | AT | AG | AC | TA | TT | TG | TC | GA | GT | GG | GC | CA | CT | CG | CC |
| --- | --- | --- | --- | --- | --- | --- | --- | --- | --- | --- | --- | --- | --- | --- | --- | --- |
| AS-AS | 4.40 | 6.20 | 3.40 | 5.20 | 2.50 | 4.40 | 1.40 | 3.30 | 3.30 | 5.20 | 2.40 | 4.20 | 1.40 | 3.40 | 0.66 | 2.40 |
| SA-SA | 4.40 | 2.50 | 3.30 | 1.40 | 6.20 | 4.40 | 5.20 | 3.40 | 3.40 | 1.40 | 2.40 | 0.66 | 5.20 | 3.30 | 4.20 | 2.40 |
| AS-SA | 6.20 | 6.20 | 5.20 | 5.20 | 6.20 | 6.20 | 5.20 | 5.20 | 5.20 | 5.20 | 4.00 | 4.00 | 5.20 | 5.20 | 4.00 | 4.00 |
| SA-AS | 6.20 | 6.20 | 5.20 | 5.20 | 6.20 | 6.20 | 5.20 | 5.20 | 5.20 | 5.20 | 4.00 | 4.00 | 5.20 | 5.20 | 4.00 | 4.00 |

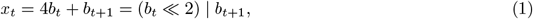

where ≪ denotes the left bitwise shift operator and | denotes the bitwise OR operator. The total energy for a window of *d* positions is

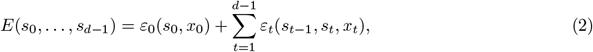

where *ε*_0_(*s*_0_, *x*_0_) is the initial-position energy term, and for *t* ≥ 1, *ε*_*t*_(*s*_*t*−1_, *s*_*t*_, *x*_*t*_) is the transition energy from the previous state to the current state at position *t*. These terms come from the original dinucleotide tables reproduced in Table 2. Brute-force evaluates all 2^*d*^ assignments of *s*_0_, …, *s*_*d*−1_. Our objective is to recover the same minimum without exhaustive enumeration.

### Canonical dinucleotide energy table

The dynamic program uses the same four transition types as the original Z-Hunt formulation [16]. Table 2 shows the canonical coefficients for the standard A/C/G/T bases as they appear in our Python code. The full implementation expands them to additional encoded symbols, but the 4 × 16 block shown here is the core model.

At runtime, the previous and current states choose a row and the observed dinucleotide chooses a column. Fig 1 shows the same model as a weighted two-state graph.

**Fig 1.**
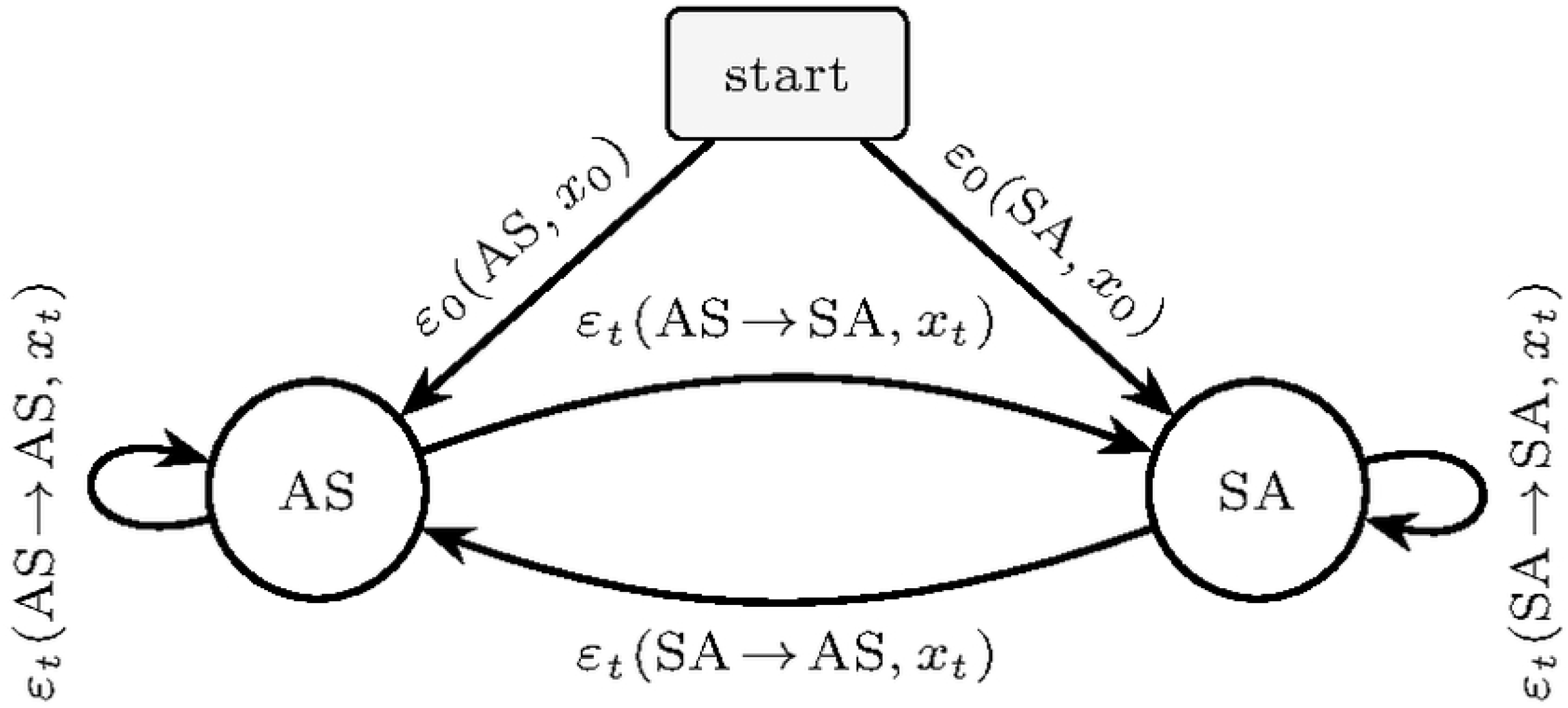
Weighted two-state representation of the canonical Z-Hunt transition table. The four transition classes in Table 2 become edges between `AS`and `SA`. The first dinucleotide contributes the two start costs depending on its state.

### Dynamic programming formulation

Dynamic programming is an optimization technique involving the decomposition of a problem into overlapping sub-problems. By solving these sub-problems and storing their results, the global solution can be derived more efficiently [22–24]. In our approach, we used a non-recursive DP implementation; this eliminates the extra memory overhead inherent to recursion, thereby guaranteeing *O*(1) space. Since Eq (2) contains only pairwise consecutive-state terms, the minimum up to position *t* depends only on the minimum ending in each state at *t* − 1. Hence, we track *DP* [*t*, AS] and *DP* [*t*, SA] which represent the minimum total energies for positions 0 through *t* ending in each state. The initialization is:

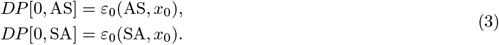

Other than the initial step, for every position *t* ≥ 1, there are exactly two ways to arrive in each state (either the previous position was `AS`or it was `SA`), so we keep the lower-energy one:

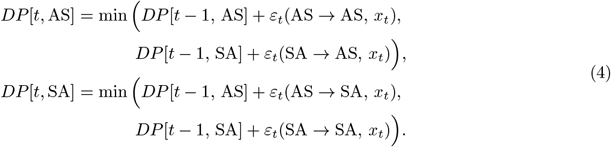

At position *d* − 1, the smaller terminal value is the minimum window energy. Stored predecessor choices recover the optimal `AS`/`SA`string and the Boltzmann-weighted quantities used later in scoring.

This reformulation reduces conformation search from *O*(2^*d*^) to *O* (*d*) time with *O*(*d*) memory for backpointers. The remaining thermodynamic calculations are already polynomial.

#### Algorithm 1

Dynamic programming optimization of anti/syn states for one window

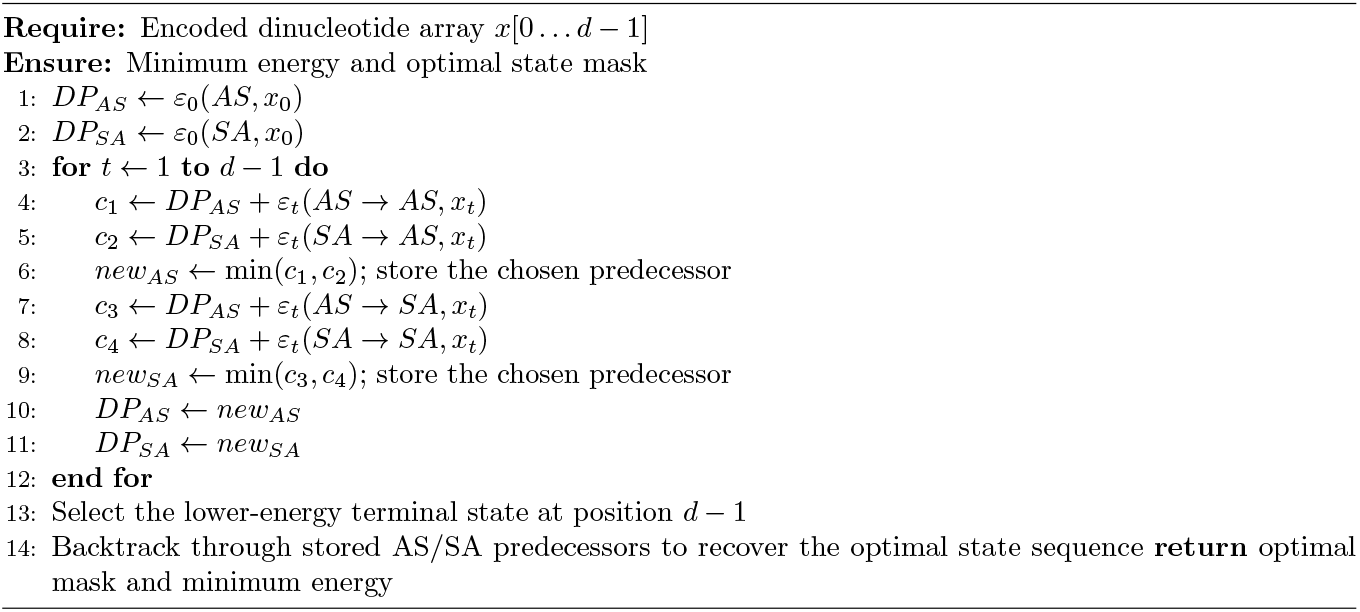

### Correctness and complexity

The recurrence above is exact because the Z-Hunt window energy decomposes into a start term plus pairwise consecutive-state terms. This is the same idea as shortest-path dynamic programming on layered graphs where at each position, we keep only the best partial path, using fixed legacy Z-Hunt dinucleotide energies as costs.

To explain exactness directly, define *F*_*t*_(*q*) as the minimum prefix energy through position *t* that ends in state *q* ∈ {AS, SA}. At *t*=0, this is exactly the initialization: *F*_0_(AS) = *ε*_0_(AS, *x*_0_) and *F*_0_(SA) = *ε*_0_(SA, *x*_0_). For *t* ≥ 1, an optimal prefix ending in *q* can only come from one of the two states at *t* − 1, so the best value at *t* is the smaller of those two predecessor choices plus the corresponding transition cost from Eq (2). This is precisely the DP recurrence, so each table entry equals the true optimum for that prefix and terminal state. Therefore, the smaller terminal value at *d* − 1 is the global minimum over all AS/SA strings. Backtracking follows through because each stored predecessor is the state that achieved the local minimum used to fill the table. Following those pointers from the chosen terminal state reconstructs an optimal path. In case ties occur, any tied predecessor chain is still optimal.

The complexity follows immediately. Each new dinucleotide position evaluates four candidate transitions and keeps two minima, so the optimization runs in *O*(*d*) time. Since the implementation is iterative, it uses only *O*(1) memory when we need the minimum energy alone, and *O*(*d*) memory when predecessor pointers are stored for path recovery.

### Python implementation and workflow

Z-Hunt-DP is a Python 3 pipeline that follows the original Z-Hunt flow. Input DNA is converted into dinucleotide indices for direct array lookup, the dynamic program is run for each window, and the inherited linking-difference statistics are then computed from the optimal state path. We speed up the core calculations with Numba, a Python library that compiles numerical functions into fast machine code at runtime, and run different windows in parallel across CPU cores using `ProcessPoolExecutor`. The pipeline also writes the raw `.Z-SCORE`file, summary tables as well as programmatically generates annotations and figures. The source code is publicly available at: https://github.com/Aljumaily/Z-Hunt-DP.

## Results

The results support two claims with different baselines. First, the dynamic program is an exact reformulation of the same optimization solved by exhaustive search, as explained in the Materials and methods section. The brute-force benchmark then provides implementation-level validation of that exactness and quantifies the runtime gain. Second, against heterogeneous public tools, we evaluate practical interval localization rather than direct thermodynamic score equivalence. Benchmarks were run on Python 3.12.10 on an Apple MacBook Air M4 with 24 GB memory.

Across all tested windows, the dynamic program matched the brute-force minimum energy within numerical tolerance. Brute-force rises from 0.0052 ms at *d*=4 to 1125.0234 ms at *d*=20, where it evaluates 2^20^ = 1,048,576 assignments, whereas dynamic programming stays between 0.000883 ms and 0.003405 ms; the speedup at *d*=20 is 3.30 × 10^5^ as depicted in Table 3 and Fig 2. This exponential divergence is expected. Brute force scales as *O*(2^*d*^) in the number of anti/syn assignments, whereas the dynamic program reduces this to *O*(*d*). Each new position evaluates exactly four candidate transitions and keeps only two running minima, so the cost of each position remains constant regardless of window length.

**Table 3.** Representative runtime comparison for one sequence window.

| $d$ | States | Brute (ms) | DP (ms) | Speedup |
| --- | --- | --- | --- | --- |
| 4 | 16 | 0.0052 | 0.000883 | $5.9\times$ |
| 8 | 256 | 0.1177 | 0.001503 | $78.3\times$ |
| 12 | 4096 | 2.7754 | 0.002209 | $1256.2\times$ |
| 16 | 65536 | 58.4407 | 0.002747 | $21271.4\times$ |
| 20 | 1048576 | 1125.0234 | 0.003405 | $330392.2\times$ |

**Fig 2.**
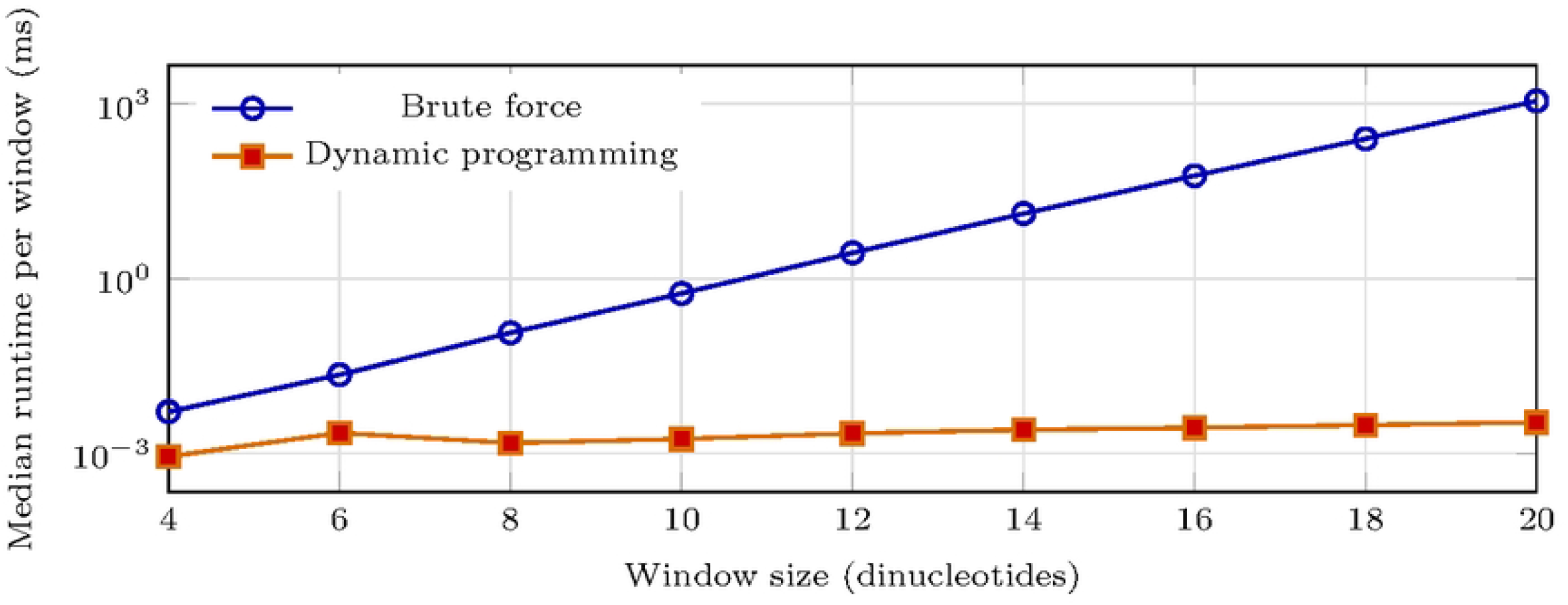
Measured per-window runtime for exhaustive search and dynamic programming from 4 to 20 dinucleotides on a logarithmic y-axis.

### Public multi-tool benchmark on experimentally mapped human loci

Since only legacy Z-Hunt solves the same objective, the broader comparison below uses interval-overlap statistics and does not imply direct thermodynamic score comparability. To evaluate practical localization, we benchmarked Z-Hunt-DP against Z-DNA Hunter, Z-DNABERT2, and ZSeeker on HeLa loci experimentally mapped using Zaa, a tandem dimer of two Z*α* domains from the Z-DNA-binding domain of human double-stranded RNA adenosine deaminase (hADAR1) [25]. Benchmarking used the six highest-signal peaks from the Zaa ChIP-seq bedGraph (GEO accession GSE71682, sample GSM1843184, hg18 assembly [26]): *ADAR*, elongation factor 1-alpha 1 (*EEF1A1*), microRNA 663 host gene (*MIR663AHG*), (hg18 chr16:33,870,062-33,870,963), (hg18 chr19:32,423,842-32,427,534), and (hg18 chr6:58,884,689-58,885,788), across four centred window lengths (24 cases). We did not rerun DeepZ because the public release lacks a packaged inference workflow, so we used the published hg19 DeepZ interval track (DeepZ.bed) only as a reproducibility reference. After converting our hg18 loci to hg19 coordinates with UCSC LiftOver, thresholding at 0.343 reproduced that published track within 14 bp in all cases. We report clipped whole-window overlap and stricter 128 bp midpoint-core localization, using covered-base Jaccard as the primary statistic:

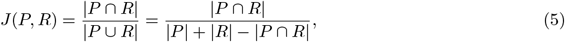

and base-level recall

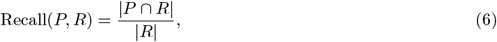

for predicted bases *P* and reference bases *R*. Aggregate statistics are the arithmetic mean over all *N* cases:

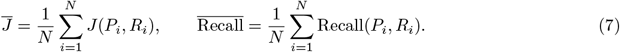

Table 4 shows that Z-Hunt-DP was the only tool to recover all 24 benchmark windows and had the strongest whole-window overlap (mean recall 1.000, mean Jaccard 0.886). Among directly rerunnable external baselines, Z-DNABERT2 was strongest on this whole-window (lenient) view, where any prediction overlapping anywhere in the evaluation window receives credit regardless of its position within that window; however, it completed only 9 of 24 cases, as the remaining 15 exceeded the 300 s service timeout of the DNA Analyser backend. Its recall and Jaccard figures are therefore computed over the nine successful cases only. ZSeeker and Z-DNA Hunter were sparse. Since several GSE71682 intervals are wider than the 128–512 bp evaluation windows, this whole-window metric is intentionally broad. DeepZ also showed broader overlap, but only as an annotation-backed reproducibility check. We used the published hg19 DeepZ.bed interval track as a static reference against which to validate our hg18-to-hg19 converted loci, rather than running DeepZ inference ourselves, since the public release provides no packaged inference workflow for arbitrary sequences.

**Table 4.** Aggregate performance on the public hg18 HeLa Zaa panel (24 cases). For service-backed tools, Req., Sleep, and Residual denote active request time, polling sleep, and remaining wall time; local tools report N/A.

| Tool | Cases | Total (s) | Req. (s) | Sleep (s) | Residual (s) | Mean recall | Mean Jaccard | Hit rate |
| --- | --- | --- | --- | --- | --- | --- | --- | --- |
| Z-Hunt-DP | 24/24 | 1.559 | N/A | N/A | N/A | **1.000** | **0.886** | **1.000** |
| ZSeeker | 24/24 | 0.794 | N/A | N/A | N/A | 0.053 | 0.053 | 0.250 |
| Z-DNABERT2 | 9/24 | 6.584 | 3.908 | 2.667 | 0.009 | 0.818 | 0.815 | **1.000** |
| Z-DNA Hunter | 24/24 | 4.798 | 3.605 | 1.188 | 0.005 | 0.001 | 0.001 | 0.083 |
| DeepZ | 24/24 | 0.000* | N/A | N/A | N/A | 0.401 | 0.401 | 0.708 |

\* Nonzero time below 0.001 s. For DeepZ, the marked value reflects liftover plus DeepZ.bed import rather than model inference.

Fig 3 shows why centre-aware evaluation matters. For MIR663AHG at 1024 bp, Z-DNABERT2 still reaches whole-window Jaccard 0.847, yet its 128 bp midpoint-core Jaccard falls to 0.051 because the merged calls leave a central gap; ZSeeker, Z-DNA Hunter, and DeepZ miss the midpoint core entirely. Midpoint-core and midpoint-anchor analyses therefore better match the follow-up task of narrowing a broad experimental interval. In practical terms, a broad overlap score can still direct follow-up assays toward the wrong part of the locus when the predicted interval is off-centre.

**Fig 3.**
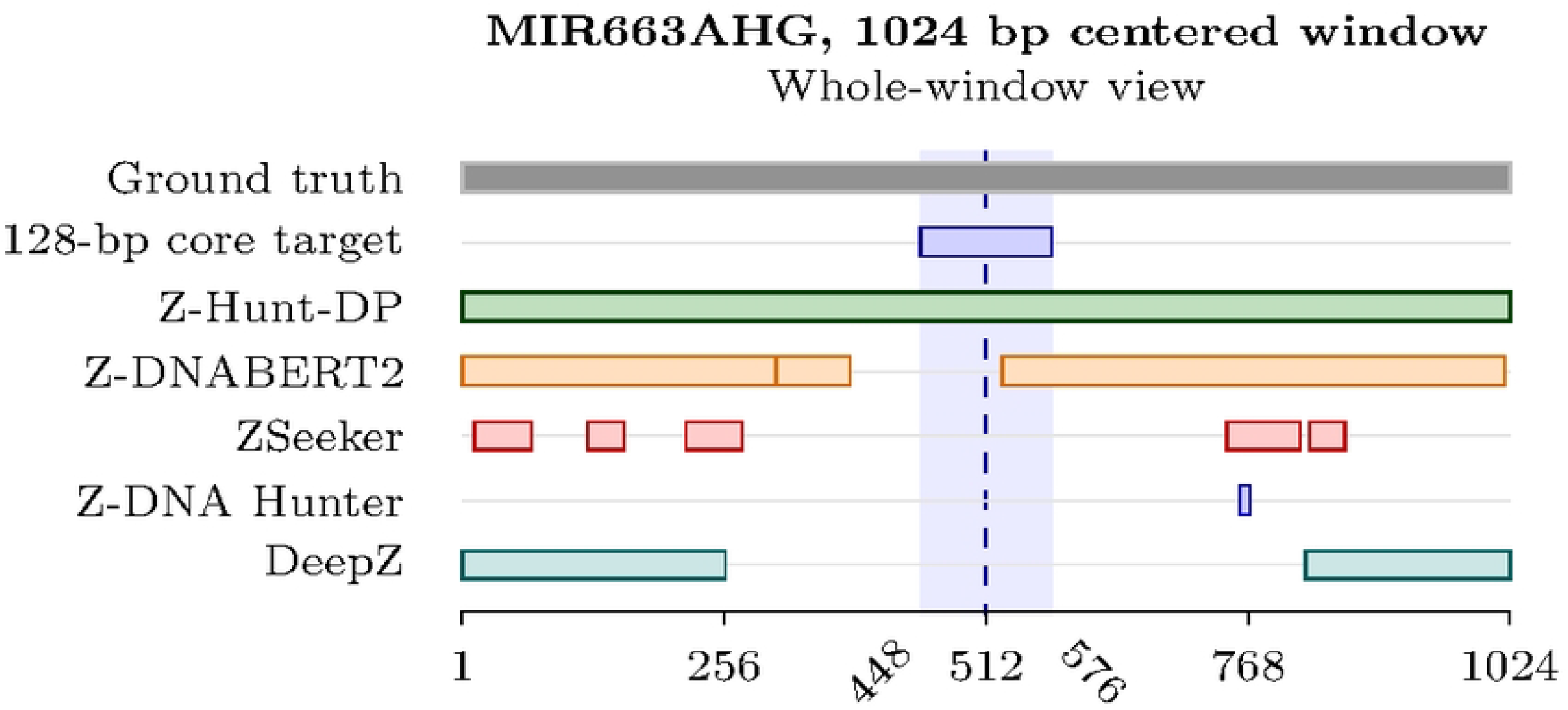
Case study at MIR663AHG in a centred 1024 bp window. The clipped target spans the full window.

Fig 4 gives the stricter midpoint-core view. Z-Hunt-DP had the highest score at every window length and maintained complete midpoint-core recall, although added flanking sequence lowered its core Jaccard by widening predicted intervals. Z-DNABERT2 was the only external tool with nonzero localization at all four lengths. ZSeeker localized only occasionally and Z-DNA Hunter remained at zero.

**Fig 4.**
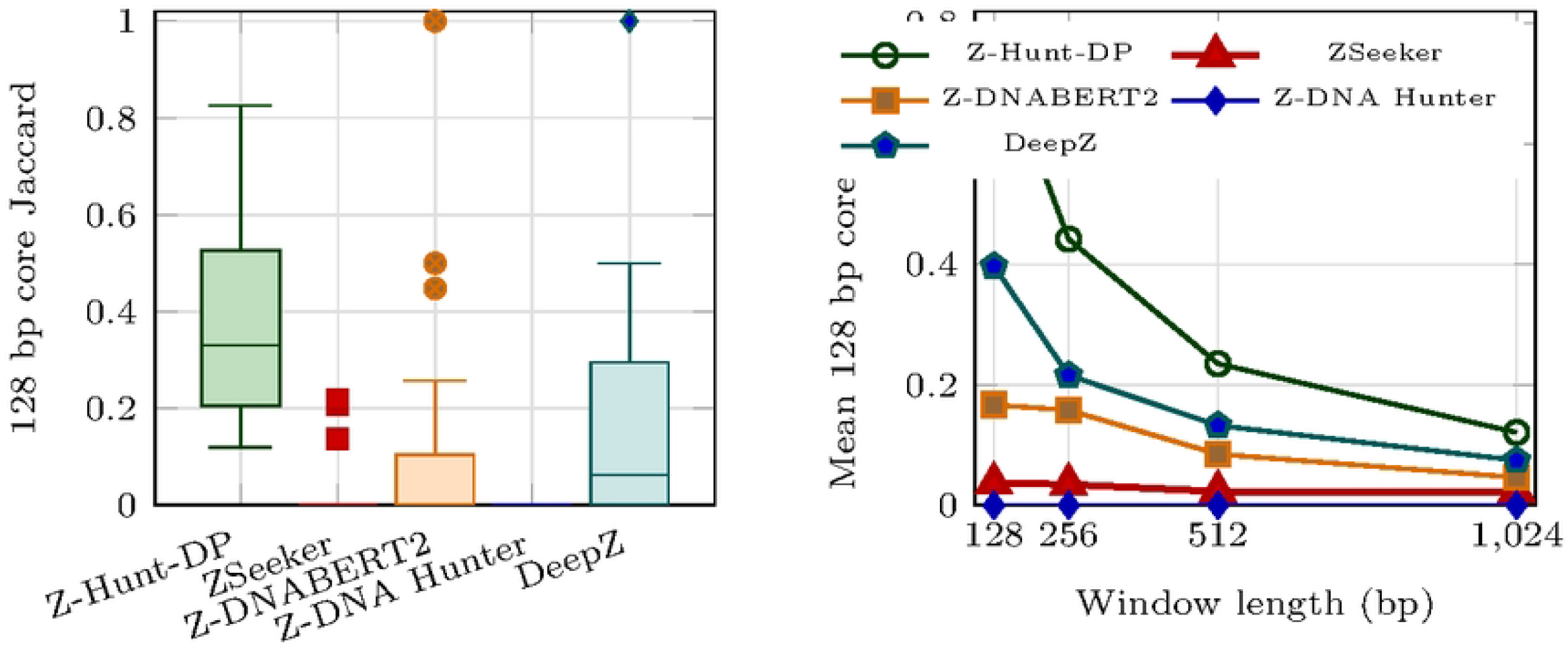
Midpoint-core localization on six hg18 HeLa Zaa loci (24 cases). Left: 128 bp core Jaccard distribution showing Z-Hunt-DP with robust signal; ZSeeker and Z-DNA Hunter show minimal to near-zero signal. Right: mean core Jaccard by window length.

We also reran the pipeline on the top 24 merged HeLa loci from the same dataset (96 cases). The ordering did not change, with Z-Hunt-DP remaining the most stable whole-window and midpoint-core localizer. As a separate throughput check on the authentic 1.50 Mb FHIT fragile-site locus (hg38 chr3:59,747,277–61,251,452), all rerunnable tools completed. Z-Hunt-DP, ZSeeker, Z-DNA Hunter, and Z-DNABERT2 finished in 231.3, 2.35, 10.8, and 251.8 seconds, respectively. Since FHIT was not part of the clipped experimental panel, we report it only as a practical scalability result.

### Additional localization stress tests

Across the same 24 cases, Z-Hunt-DP was also the only method with perfect hit rate when the midpoint core was tightened from 128 bp to 16 bp and the only one to hit the exact midpoint anchor at every length. DeepZ and Z-DNABERT2 retained partial signal, hitting 8 and 6 of 24 exact anchors, whereas ZSeeker recovered midpoint-core overlap only at 64 bp and Z-DNA Hunter remained at zero.

Miss distances show that these failures were often substantial rather than near-boundary ambiguities. Among miss-only detections, the median nearest-anchor distance was 11 bp for Z-DNABERT2, 29 bp for ZSeeker, 105 bp for DeepZ, and 309 bp for Z-DNA Hunter. The stress tests therefore distinguish tools that remain centered from tools that merely land somewhere inside a broad experimental interval.

That robustness is consistent with the case study view. Z-Hunt-DP continues to prioritize the midpoint itself rather than only a flanking subset of the broader experimental interval.

## Conclusion and future directions

Z-Hunt-DP exactly reformulates the original Z-Hunt conformation optimization problem, preserving the legacy thermodynamic objective while replacing exponential anti/syn search with linear-time optimization. On tested windows it matched brute-force minimum energies and achieved speedups up to 3.30 × 10^5^. Under interval-based evaluation against heterogeneous external tools, it also remained the strongest and most stable localizer across whole-window overlap, midpoint-core overlap, exact-midpoint stress tests, and expanded-panel reruns. These cross-tool results remain interval-based rather than direct thermodynamic score comparisons, DeepZ remains an annotation-backed reproducibility check, and service-backed tools may change under different deployments. Within those limits, Z-Hunt-DP provides an auditable open-source route to exact optimization under the legacy Z-Hunt model. For practical follow-up studies, the important point is that this advantage is not only broader interval recovery but more reliable centering on the experimentally supported locus. This pattern persisted under narrower midpoint cores, exact-anchor tests, and expanded-panel reruns, making the method useful both for broad screening and for prioritizing short validation constructs around a nominated site. Since the optimization remains exact and inspectable, downstream interval calls can be traced back to a transparent thermodynamic objective rather than a hidden model state or a service revision. This distinction matters most when experimentally mapped intervals are wide. A method can appear concordant at the whole-window level while still shifting the candidate site away from the biologically relevant center. In that setting, exact dynamic programming provides not only speed and reproducibility but a better starting point for focused perturbation, validation, and sequence-design follow-up.

Future work will expand the supported bases to include 5-methylcytosine (5mC), 5-hydroxymethylcytosine (5hmC), 5-formylcytosine (5fC), and 5-carboxylcytosine (5caC). These non-cannonical nucleotides occur as epigenetic modifications whose effect on the B-to-Z transition energy is shown in both computational and physical chemistry literature [27]. Moreover, a natural extension of this work would be to augment the dynamic programming framework with reinforcement learning style updates, allowing local decisions to be refined progressively from sequence-level feedback rather than being fixed only after a complete global pass.

## Data availability

The benchmark data and all scripts used to reproduce the results in this article are available at https://github.com/Aljumaily/Z-Hunt-DP.

## Author contributions

M.A.: methodology, implementation, experiments, writing - drafting. H.Q.: data collection & curation, result validation. H.Y.: conceptualization, supervision, writing - editing. Y.L.: supervision, resources, writing - editing.

## Acknowledgments

We wish to thank the Shared Hierarchical Academic Research Computing Network (SHARCNET) and Digital Research Alliance of Canada for providing access to computational facilities at the early stage of this work.

